# The zebrafish meiotic cohesion complex protein Smc1b is required for key events in meiotic prophase I

**DOI:** 10.1101/2021.05.25.445674

**Authors:** Kazi Nazrul Islam, Maitri Mitesh Modi, Kellee Renee Siegfried

## Abstract

The eukaryotic structural maintenance of chromosomes (SMC) proteins are involved in key processes of chromosome structure and dynamics. SMC1β was identified as a component of the meiotic cohesion complex in vertebrates, which aids in keeping sister chromatids together prior to segregation in meiosis II and is involved in association of homologous chromosomes in meiosis I. The role of SMC1β in meiosis has primarily been studied in mice, where mutant male and female mice are infertile due to germ cell arrest at pachytene and metaphase II stages, respectively. Here, we investigate the function of zebrafish Smc1b to understand the role of this protein more broadly in vertebrates. We found that zebrafish *smc1b* is necessary for fertility and has important roles in meiosis, yet has no other apparent roles in development. Therefore, *smc1b* functions primarily in meiosis in both fish and mammals. In zebrafish, we showed that *smc1b* mutant spermatocytes initiated telomere clustering in leptotene, but failed to complete this process and progress into zygotene. Furthermore, mutant spermatocytes displayed a complete failure of homolog pairing and synapsis. Interestingly, meiotic DNA double strand breaks occurred in the absence of Smc1b despite failed pairing and synapsis. Overall, our findings point to an essential role of Smc1b in the leptotene to zygotene transition during zebrafish spermatogenesis. In addition, ovarian follicles failed to form in *smc1b* mutants, suggesting an essential role in female meiosis as well. Our results indicate that there are some key differences in Smc1b requirement in meiosis among vertebrates: while Smc1b is not required for homologue pairing and synapsis in mice, it is essential for these processes in zebrafish.

## 1 INTRODUCTION

Germ cells are unique since only they can undergo meiosis and contribute to the next generation by producing haploid gametes. Meiosis is a specialized cell division where DNA replicates once but chromosomes go through two rounds of segregation (Bolcun-Filas and Handel, 2018). The first meiotic division (meiosis-I) segregates homologous chromosome pairs whereas the second division (meiosis-II) is more similar to mitotic divisions, separating sister chromatids. To accomplish the unique segregation in meiosis-I, homologous chromosomes (each of which consists of two sister chromatids) must find each other, pair, and undergo synapsis. In many eukaryotic organisms, such dynamic chromosome movements that assist in homologue pairing are facilitated by telomeres (Alleva and Smolikove, 2017). Telomeres attach to the nuclear envelope during the leptotene stage via a meiotic-specific protein complex. This complex interacts with a protein chain that spans the nuclear envelope and interacts with the cytoskeleton to drive chromosome movements (Burke, 2018). The telomeres then cluster at one side of the nucleus, forming the “bouquet”. This process of chromosome movement and clustering is thought to be essential for homologous chromosomes to find each other and pair.

As homologous chromosomes pair, a proteinaceous structure assembles between them, called the synaptonemal complex (SC), which holds homologous chromosomes together and facilitates meiotic recombination (Bolcun-Filas and Handel, 2018). The vertebrate SC consists of 3 synaptonemal complex proteins (SYCP): SYPC2 and SYCP3 form the axial elements, which associates with the chromosome axes; and SYPC1 forms the transverse elements, which bridges the axial elements of homologous chromosomes. In addition, multiple SC central element proteins overlap with the transverse element as chromosomes synapse. The formation of the synaptonemal complex between homologous chromosomes is critical for segregation of homologous chromosomes to opposite daughter cells during the first meiotic division. In zebrafish, SC formation is initiated near the telomeres (Saito et al., 2014; Blokhina et al., 2019). The axial element of the SC assembles along chromosomes beginning near the chromosome ends, as has been visualized by Sycp2 and Sycp3 localization (Saito et al., 2011; Blokhina et al., 2018; Takemoto et al., 2020). Synapsis ensues, as seen by visualization of the transverse element protein, Sycp1, which follows Sycp3 localization (Blokhina et al., 2018; Imai et al., 2021). Thus, homologue pairing and synapsis begins at the chromosomes ends and zippers closed towards the chromosome center. As homologue pairing commences, meiotic double strand breaks, which are a prerequisite for homologous recombination, also initiate near the chromosome ends in zebrafish (Saito et al., 2011; Blokhina et al., 2018).

Formation of the SC in meiosis is dependent on the cohesion complex. The cohesion complex has a critical role in the faithful pairing and segregation of chromosomes during both mitosis and meiosis (Ishiguro, 2019). In both processes the cohesion complex promotes sister chromatid cohesion and proper chromosome segregation, however in meiosis it has additional roles in homologue pairing, assembly of the synaptonemal complex, and chromosome architecture. The meiotic cohesion complex consist of four core subunits: SMC1β, SMC3, RAD21L or REC8, and STAG3 (Fig 1A) (Rankin, 2015). Similarly, mitotic cohesion consists of SMC1α, SMC3, RAD21, and STAG1/2 (Nasmyth, 2001). The two SMC proteins form a ring surrounding the sister chromatids that is closed at one end by RAD21L/REC8 and STAG3 (or RAD21 and STAG1/2 in mitosis). This complex holds the sister chromatids together until anaphase, when separase cleaves RAD21L/REC8/RAD21, allowing sister chromosomes to move to opposite poles of the cell (Rankin, 2015). In meiosis, the cohesion complex is necessary for assembly of the SC axial element, and therefore promotes synaptonemal complex formation. The specific components of each cohesion complex contributes to their unique roles in meiosis and mitosis (Ishiguro, 2019).

**Figure 1:**
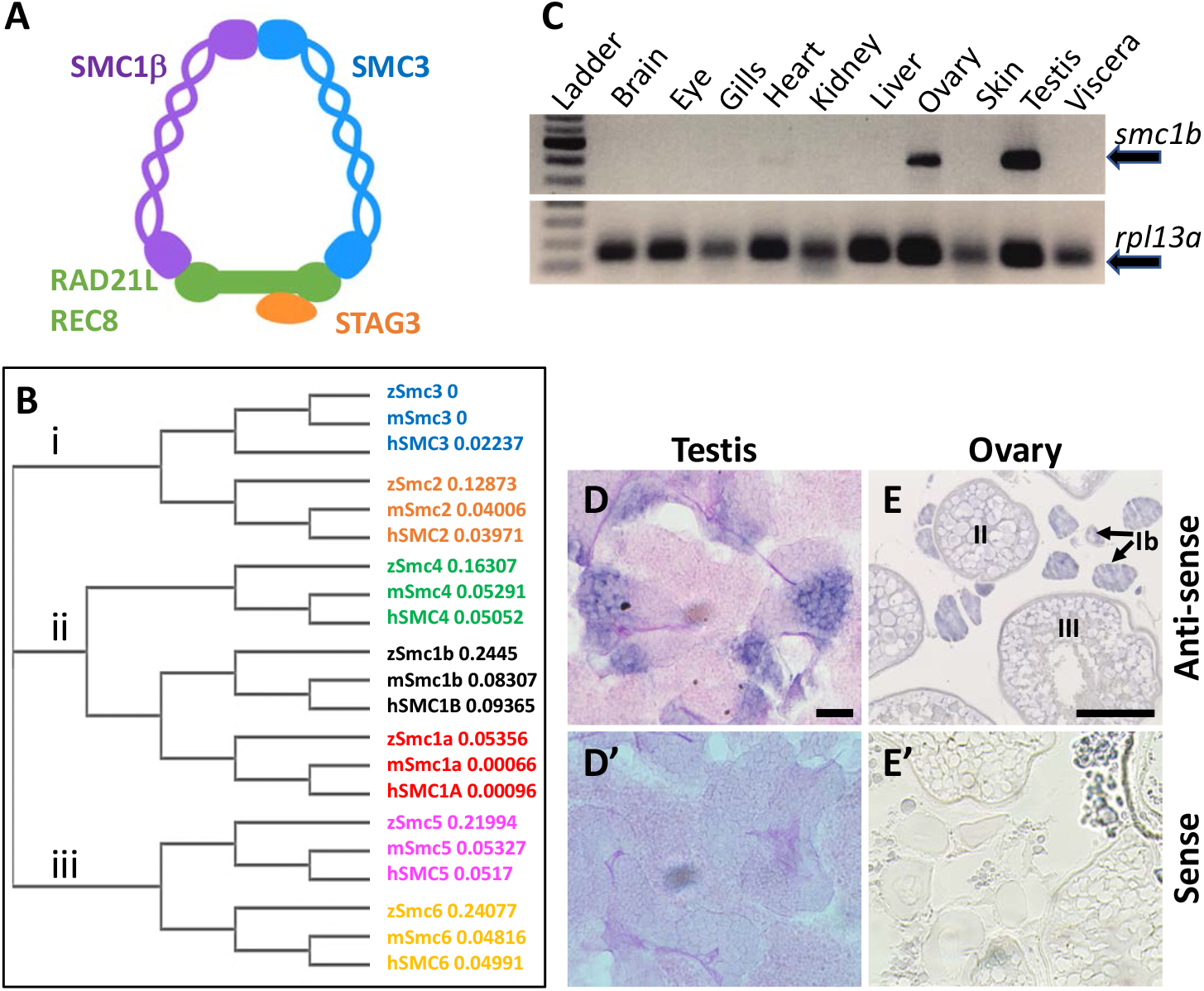
Zebrafish *smc1b* is conserved and expressed in germ cells. (A) Cartoon representing the meiotic cohesion complex. (B) Phylogenetic tree of zebrafish (z), mouse (m), and human (h) Smc protein family. Numbers to the right of protein names represent bootstrap confidence values. (C) RT-PCR of *smc1b* and control gene *rpl13a* using cDNA generated from brains, eyes, gills, hearts, kidneys, livers, ovaries, skin, testes, and viscera of wild-type adult zebrafish. (D-E) *In-situ* hybridization (ISH) on testis (D) and ovary (E) sections of wild-type zebrafish. ISH using sense probes were run as negative controls. Scale bars are 20 μm for testes and 100 μm for ovaries.

Vertebrate animals typically have two SMC1 proteins, the mitotic SMC1α and meiotic SMC1β, however the distinct roles of these two proteins have not been studied outside of mammals. Mice lacking SMC1β fail to complete meiosis and present as infertile in both sexes, indicating that it is specifically necessary for meiosis (Revenkova et al., 2004; Takabayashi et al., 2009). In *smc1β* mutant mice, meiosis is blocked in pachytene in males whereas female meiosis progresses up to metaphase II. The other observed defects in the mutant mice were short prophase-I axial elements, incomplete synapsis, premature loss of sister chromatid cohesion, and reduced recombination foci (Revenkova et al., 2004). In addition, mammalian SMC1β has a role in telomere integrity and attachment of telomeres to the nuclear envelope during meiotic prophase-I. In mice, SMC1β is required for localization and enrichment of cohesion to telomeres in meiotic prophase-I and *smc1β* deficient cells display various types of telomere abnormalities (Adelfalk et al., 2009). In addition, around 20% of telomeres in the *smc1β* mutant meiocytes failed to attach to the nuclear envelope during meiosis. Thus, this protein is necessary for multiple processes required for proper chromosome segregation during meiosis.

Whether or not SMC1β functions in meiosis in non-mammalian vertebrates has not been investigated, although it was found to be expressed in medaka spermatocytes (Iwai et al., 2004). Here, we analyzed the function of zebrafish *smc1β,* which is named *smc1b* in zebrafish. We found that all *smc1b* mutant zebrafish develop as males with no other outward visible defects. Histological examination of the testes revealed that they failed to develop mature sperm and consequently were sterile. *Smc1b* mutant spermatocytes failed to progress through meiotic prophase-I, demonstrating that *Smc1b* is specifically necessary for meiosis in zebrafish, similar to mice. Interestingly, zebrafish *smc1b* mutant males exhibited more severe meiotic defects than those reported in mice, displaying meiotic arrest at the leptotene stage, whereas mouse spermatocytes reached up to the pachytene stage. Furthermore, the cohesion complex did not appear to be enriched at telomeres in zebrafish spermatocytes, however meiotic bouquet formation was still disrupted in mutants. This study broadens our understanding of how the meiotic cohesion complex functions in vertebrate animals.

## 2 MATERIALS AND METHODS

### 2.1 Zebrafish strains and husbandry

All animal procedures were approved by the UMass Boston Animal Care and Use Committee. Zebrafish were maintained in a recirculating system on a 14hr light and 10hr dark light cycle. Strains and lines used in this study were: *smc1b^sa24632^, Smc1b^sa24631^*, and Tuebingen (Tue). Embryos of two *smc1b* mutant lines (ID: sa24632 and sa24631) were generated by the Zebrafish Mutation Project and obtained from the Zebrafish International Resource Center (ZIRC). The *smc1b*^sa24632^ and *smc1b*^sa24631^ lines each carry a nonsense mutation which cause premature stop codons at codon 198 and 261, respectively, whereas the wild-type protein is 1235 amino acids long. For simplicity, we refer to each mutant line as *smc1b^Q198X^* and *smc1b^Q261X^*.

### 2.2 Protein alignment

We used the online program MUSCLE [https://www.ebi.ac.uk/Tools/msa/muscle/] to perform protein alignment and phylogenetic tree generation. The NCBI accession number of the proteins used in this study are as follows: zebrafish Smc1a (NP_001155103.1), Smc1b (XP_009296271.1), Smc2 (NP_955836.2), Smc3 (NP_999854.1), Smc4 (NP_775360.2), Smc5 (NP_001180470.1), and Smc6 (NP_001121806.1); mouse Smc1a (NP_062684.2), Smc1b (NP_536718.1), Smc2 (NP_001288341.1), Smc3 (NP_999854.1), Smc4 (AAH62939.1), Smc5 (AAH38345.1), and Smc6 (AAH90630.1); human [SMC1A (CAI42646.1), SMC1B (AAI26209.1), SMC2 (AAI44164.1), SMC3 (NP_005436.1), SMC4 (Q9NTJ3.2), SMC5 (NP_055925.2), and SMC6 (CAC39248.1).

### 2.3 RT-PCR

Total RNA was isolated from zebrafish organs using TRI-reagent, following the manufacturer’s procedure. Isolated RNA was treated with TURBO DNase to remove genomic DNA contamination. The first strand cDNA was synthesized using oligo-dT primers and AMV-Reverse Transcriptase. The *ziwi* transcript, which is expressed in the germ cells of both sexes, was used as a control for the gonad using previously published primers (Siegfried and Nüsslein-Volhard, 2008). The ubiquitously expressed gene *ribosomal protein L13 alpha (rpl13a)* was used as control for other tissues (Tang et al., 2007). The primers used for RT-PCR are listed in table 1.

**Table 1:**
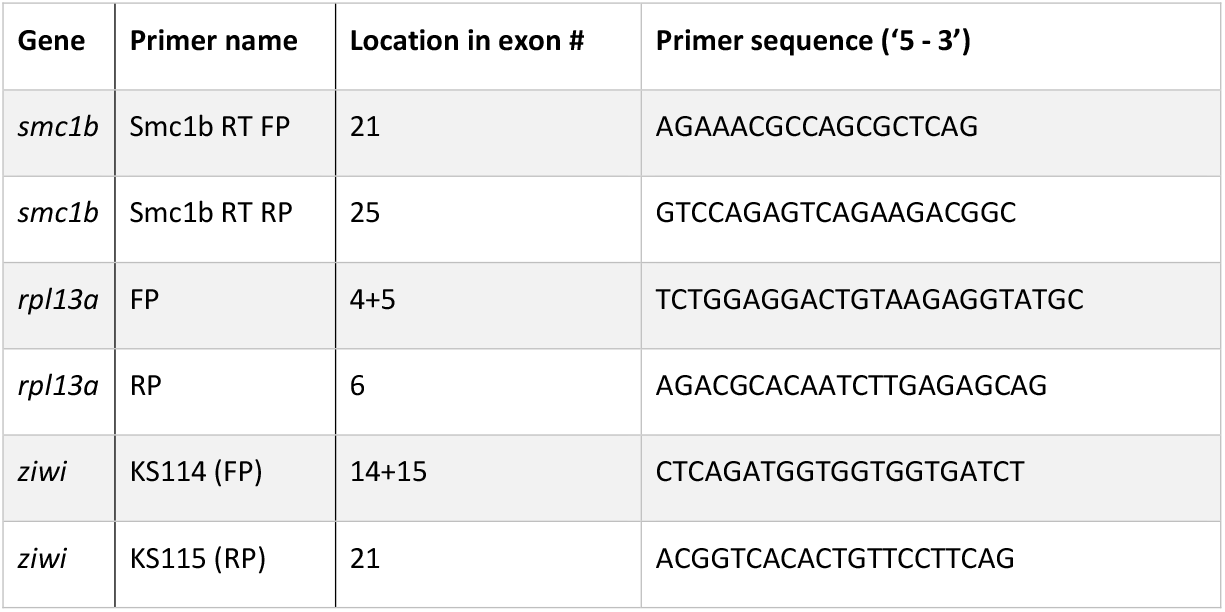
RT-PCR primers.

### 2.4 *In situ* hybridization (ISH)

Samples for ISH were fixed in 4% paraformaldehyde (PFA) at 4°C overnight. To generate a probe for detecting *smc1b,* a 390 bp long partial cDNA was made using *smc1b* RT-PCR primers (Table 1) and sub-cloned into the pGEMT Easy vector [Promega]. Both sense and anti-sense probes were synthesized using a DIG RNA Labelling kit. The ISH on paraffin-embedded sections were performed according to the protocol described by Webster *et al* (Webster et al., 2019). Alkaline phosphatase-conjugated anti-DIG antibody [Roche] and BM Purple substrate were used to visualize the probe.

### 2.5 Genotyping and PCR

To genotype the *smc1b* mutants, nested PCR reactions were performed, with the second PCR reaction using dCAPS (Derived Cleaved Amplified Polymorphic Sequences) primers, designed using the online tool dCAPS Finder 2.0 (http://biology4.wustl.edu/dcaps/). In the first PCR, primers smc1b FP and smc1b RP1 were used to amplify a 751 bp amplicon, which contained both mutant alleles. The nested PCR reaction for the *smc1b^Q198X^* line was performed using primers smc1b_sa24632 FP & RP, followed by Hpy188I digestion. The digested PCR product was run on 7% acrylamide gel to distinguish mutant (127 bp) and wild-type (152 bp) bands. Primers smc1b_sa24631 FP & RP2 were used to amplify the *smc1b^Q261X^* mutation region, followed by DdeI digestion. To distinguish the mutant (182 bp) from wild-type (157 bp) band, the digested PCR product was run on a 7% acrylamide gel. Sanger sequencing were done on PCR products to confirm all of the mutations. We also performed sequencing on cDNA from homozygous mutant testes and detected the predicted mutations. The name and sequence of the genotyping primers are listed in table 2.

**Table 2:**
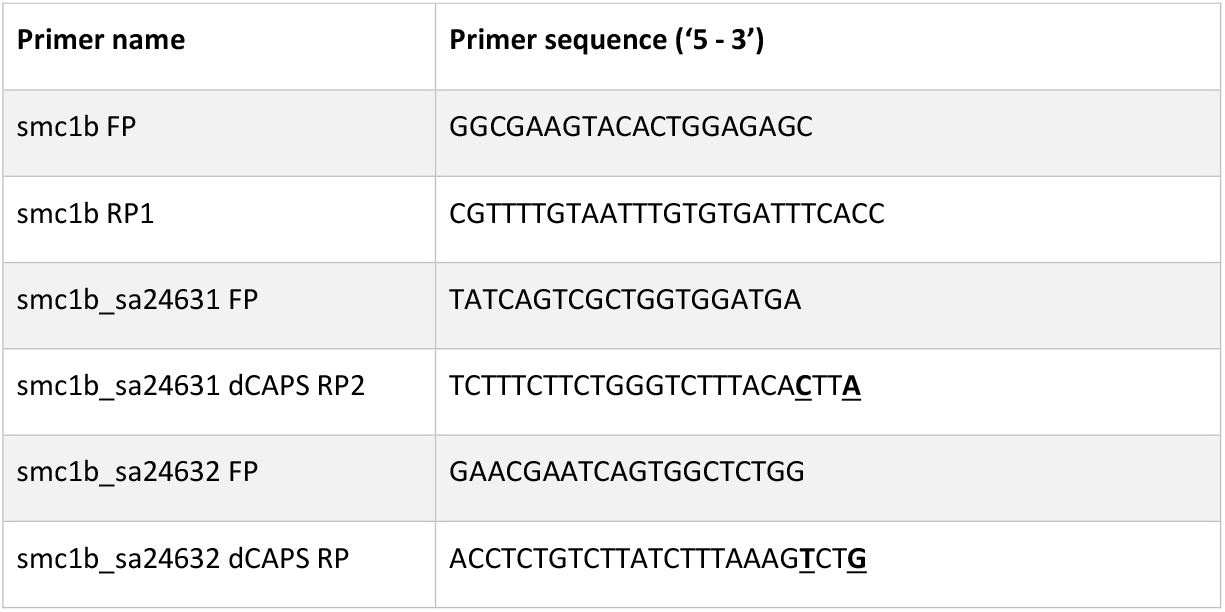
Genotyping primers. Mismatch bases are underlined

### 2.6 Fertility tests

*Smc1b^Q198X^* mutant males were paired with wild type Tue females. The following morning, embryos were collected and monitored under a dissecting microscope to see if embryonic cell cleavages took place. If egg appeared unfertilized, they were kept up to 24 hours to ensure development did not proceed. Eggs were scored as unfertilized if there was no apparent initiation of embryonic development. The embryos were raised in 1x E3 buffer in a 28°C incubator.

### 2.7 Histology

Fish were euthanized by tricaine overdose and torsos were isolated and fixed in Bouin’s solution overnight at room temperature. Following dehydration, the fixed tissues were embedding in paraffin for sectioning. A rotating microtome was used for obtaining 5 μm thick sections. We used Modified Harris’s hematoxylin and eosin for histological staining following standard protocols. Imaging was performed by a Zeiss Inverted Microscope and acquired using Zen Software.

### 2.8 Immunofluorescence and telomere staining on sections

Testes were isolated and fixed in 4% PFA for overnight at 4°C. The fixed tissues were embedded in paraffin and cut into 5 μm thick sections. After de-paraffinization and rehydration, antigen retrieval was done by heating the sections in 10 mM sodium citrate, pH 6.0 solution for 30 minutes using a vegetable steamer. After 3×5 minutes washes in PBST, the sections were circled with a barrier pen, covered with blocking buffer (1% bovine serum albumin in PBST) and incubated for 30 minutes in a humidified chamber at room temperature. Primary antibodies were diluted in the blocking buffer as follows: Sycp3 (NB300-232, Novus Biologicals) at 1:200, γ-H2AX (Neumann et al., 2011) at 1:200. The sections were incubated overnight with the primary antibodies at 4°C. Incubation with anti-rabbit Alexa Fluor Plus 488 [Thermo Fisher Scientific] secondary antibody (1:500) was done for 1 hour at room temperature. Telomeres were stained using a TelC-Cy3 probe [PNA Bio] following previously described protocol (Saito et al., 2014). To visualize nuclei, sections were counterstained with DAPI for 10 minutes and cover slipped using Fluoroshield (Sigma-Aldrich) mounting medium.

### 2.9 Chromosome spread preparation, telomere staining, and immunofluorescence

Meiotic chromosome spreads from zebrafish testes were prepared according to the previously published protocol (Blokhina et al., 2019), except that cell membranes were disrupted by incubating isolated cells in 0.8% sodium citrate solution for 20 minutes instead of 0.1 M sucrose solution. Spreads were preserved in −20°C before performing further analysis. Telo-FISH on spreads was done following the same protocol applied to sections. For antibody labelling, primary antibody Sycp1 (Blokhina et al., 2019), Sycp3 [NB300-232, Novus Biologicals], Smc3 [PA529131, Thermo Fisher Scientific], and Rad51 [GTX100469, GeneTex] were diluted at 1:200 and incubated overnight at 4°C. Incubation in appropriate secondary antibody (1:500) was done for 1 hour at room temperature. Spreads were kept at 4°C before imaging by a confocal microscopy [Zeiss LSM 880].

### 2.10 Image Analysis

Image acquisition was done with ZEN Black software attached to the confocal microscopy. Cell counting and further analysis was performed using open-sourced software Fiji/Image J. We performed Student’s t-test (p < 0.05) to see whether difference between wildtype and mutant caspase-3 positive cells were significant or not.

## 3 RESULTS

### 3.1 Zebrafish Smc1b has key domains conserved with mammals

To investigate the evolutionary conservation of Smc proteins among the vertebrates, we generated a phylogenetic tree comparing zebrafish, mouse, and human Smc protein sequences. The tree has three major branches which consist of: (i) Smc2 and Smc3; (ii) Smc1a, Smc1b, and Smc4; (iii) Smc5 and Smc6 (all SMC1α/Smc1a and SMC1β/Smc1b are referred to as Smc1a and Smc1b, respectively, for simplicity) (Fig 1B). The tree demonstrates that each zebrafish Smc protein clusters together with the corresponding mammalian protein. We also found that most mouse and human SMC proteins are evolutionary closer to each other than to those of zebrafish. Surprisingly, mouse SMC3 grouped with zebrafish Smc3 rather than that of human (Fig 1B). We focused our attention to the Smc1b protein since previous studies demonstrated a role of this protein in meiosis and reproduction (Revenkova et al., 2004). To investigate Smc1b conservation among vertebrates, we performed a protein alignment, which showed that zebrafish Smc1b is 53.04 and 52.11% identical to human and mouse SMC1β, respectively (Supplemental Fig 1). Higher similarity was observed in several key domains, such as the N-terminal ATP binding cassette (ABC), the flexible hinge, and the C-terminal ABC domains, which are 62.75, 66.95, and 75.96 % identical between zebrafish and human, respectively (Supplemental Fig 1B). However, coil-coiled domains, which are located between the ABCs and the hinge domain, are less conserved (ranging between 26.53 to 57.14%). Interestingly, we found four conserved motifs (Walker A & B, ABC transporter signature motif, and D-Loop) in zebrafish Smc1b, similar to mammals (Supplemental Fig 1) (Revenkova et al., 2001). Walker A motifs have been shown to bind with azido-ATP, an analogue of ATP (Akhmedov et al., 1998). The signature motif is essential for head-to-head engagement of SMC dimers and the Walker B sequence is involved in ATP hydrolysis (Chao et al., 2017). These results suggest that the functions of zebrafish Smc1b could be similar to the mammalian protein.

### 3.2 *Smc1b* is required for spermatogenesis and oogenesis in zebrafish

To ask which zebrafish tissues *smc1b* may function in, we assessed *smc1b* expression. We performed RT-PCR from brains, eyes, gills, hearts, kidneys, livers, ovaries, skin, testes, and viscera of wild-type adult zebrafish to test which organs of zebrafish express *smc1b*. *Smc1b* was detected in the testis and ovary but not in other organs tested (Fig 1C), suggesting that *smc1b* has important functions in zebrafish gonads. Next, we sought to answer which gonadal cells express *smc1b* by performing *in-situ* hybridization (ISH) on testis and ovary sections. The ISH detected expression in the spermatogonia and some spermatocytes but not in the mature sperm of the testis (Fig 1D). In ovaries, *smc1b* was expressed in stage Ib, II, and III oocytes, which are all in meiosis-I (Fig 1E). The ISH results bolstered the notion that *smc1b* has important roles in zebrafish germ cells.

To test whether *smc1b* is required for meiosis in zebrafish, we analyzed two mutant lines. The *smc1b^sa24632^* and *smc1b^sa24631^* mutations each result in a premature stop codon after amino acid 197 and 260, respectively. Here, we refer to these mutant lines as *smc1b^Q198X^ and smc1b^Q261X^,* respectively, reflecting the protein change caused by each mutation (Fig 2A). The homozygous mutants of both alleles developed normally without showing any outwardly visible defects. However, all *smc1b* mutant fish (N=54) developed as phenotypic males (based on pigmentation and body shape). When mutant males were paired with wild-type females, they were able to induce spawning (N=5), however all eggs were unfertilized (Table 3). To identify the underlying problem, *we* performed histology on testes from adult mutants and wild-type siblings (Fig 2). The histology revealed complete lack of spermatozoa in both mutant lines, however spermatogonia and spermatocytes were present suggesting that mutant germ cells had arrested as spermatocytes (Fig 2C-D). We also established a trans-heterozygous *(smc1b^Q198X/Q261X^*) line, which exhibited phenotypes indistinguishable to the homozygous mutants, i.e. all male and no sperm (Fig 2E). To test if *smc1b* mutant testes had abnormal cell death, we performed Immunofluorescence (IF) on mutant and wild-type testes (N=3) with cleaved caspase-3 antibody. There was no significant difference between wild-type and mutant testes in terms of apoptosis (Fig 2F-H). Therefore, *smc1b* mutant cells that arrest in meiosis die by a caspase-3 independent mechanism. These data point to a potential role of zebrafish *smc1b* in meiosis.

**Figure 2:**
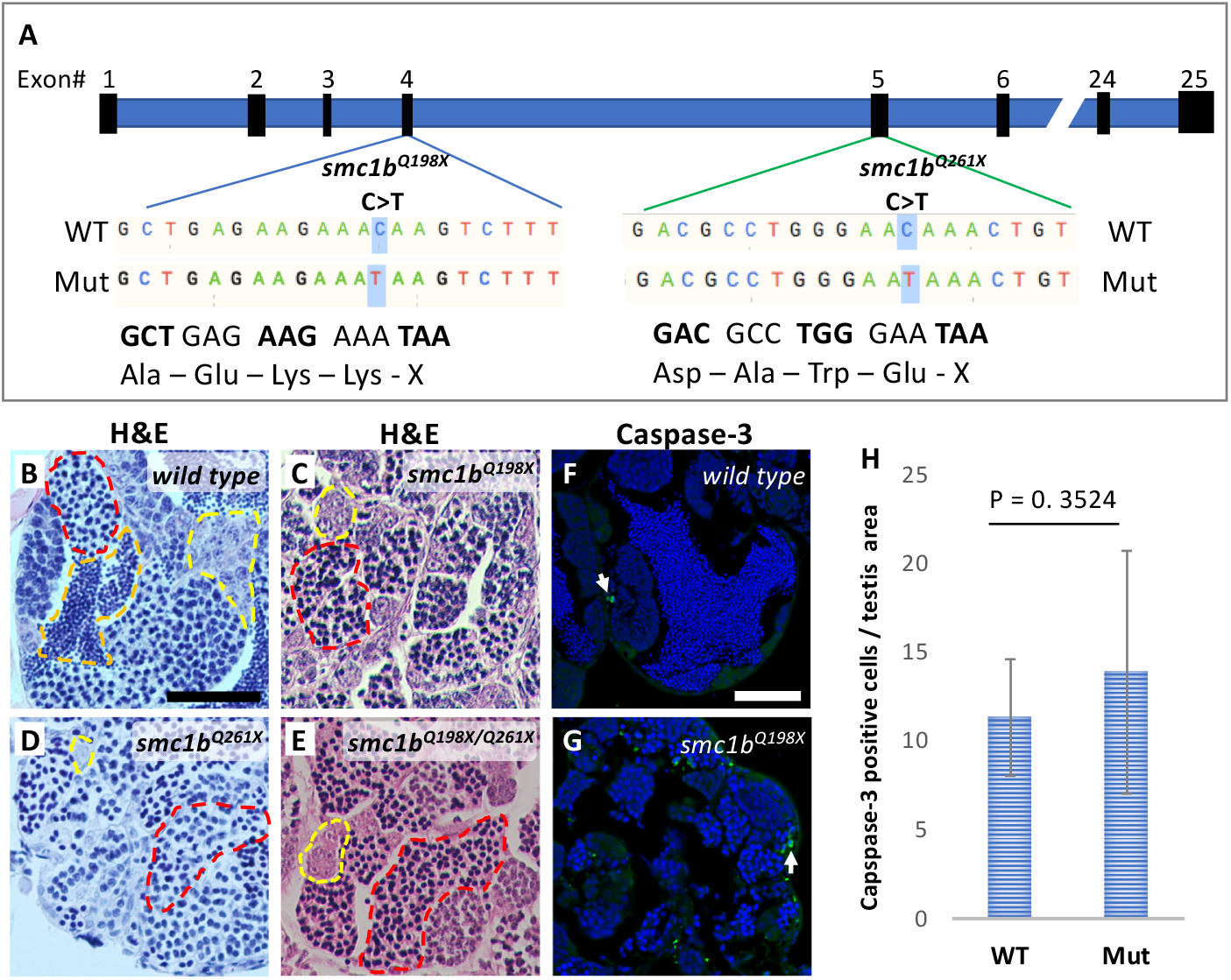
*Smc1b* mutant zebrafish display defects in spermatogenesis. (A) *Smc1b* mutations used in this study: *smc1b^Q198X^* located in exon 4 and *smc1b^Q261X^* in exon 5. Both mutations cause premature termination codons. (B-G) Hematoxylin-eosin (H&E) staining on adult testes of wild type (B), *smc1b^Q261X(-/-)^* (C),*smc1b^Q198X(-/-)^* (D), and *smc1b^Q198X/Q261X^* (E). Dashed lines denote examples of: spermatogonia (yellow), spermatocytes (red), and spermatozoa (orange). (F-G) Caspase-3 (green) and DAPI (blue) staining on adult testes from wild type (F) and *smc1b* mutant (G). Caspase-3 positive cells are indicated by arrows. (H) Quantification of caspase-3 positive cells per testis area in the mutant (N=3 testes) and wild type (N=3 testes). Student’s t-test shows no significant (p = 0.3524) difference between wild-type and mutant caspase-3 positive cells numbers. Scale bars are 50 μm.

**Table 3:** *Smc1b* mutant males are infertile. SD: standard deviation

| Genotype | Number of crosses | Numbers of eggs per cross<br>(Mean ± SD) | Percentage of fertilized eggs<br>(Mean ± SD) |
| --- | --- | --- | --- |
| *smc1b*<sup>+/+</sup> | 5 | 65.00 ± 13.21 | 97.11 ± 3.40 |
| *smc1b*<sup>Q198X</sup> | 5 | 92.6 ± 19.11 | 0.0 ± 0.0 |

Since all *smc1b* mutants were male as adults, we asked if mutants underwent sex reversal earlier in development. In zebrafish, undifferentiated gonads first pass through a “juvenile ovary” stage, in which the gonads contain immature oocytes, before undergoing sex-differentiation to form either an ovary or testis (Takahashi, 1977). Sex-differentiation of the gonads was histologically apparent at 5 weeks post fertilization (wpf) in our lines (Fig 3). To test if *smc1b* mutants generated immature oocytes and initiated female differentiation, we performed histology on mutants and wild-type siblings at 4, 5, and 6 wpf. At 4 wpf, all mutant and wild-type fish had undifferentiated gonads with no oocytes, indicating that they had not yet reached the juvenile ovary stage (Fig 3A-B). However, histology at 5 and 6 wpf revealed that 100% of *smc1b* mutant gonads were developing as testes, whereas the gonads of the wild-type siblings were either testes or ovaries (Fig 3C-H). These results indicate that *smc1b* mutants did not undergo sex reversal rather directly developed as male. Since we never detected ovarian follicles in the *smc1b* mutants, we conclude that this gene is also necessary for oogenesis in zebrafish.

**Figure 3:**
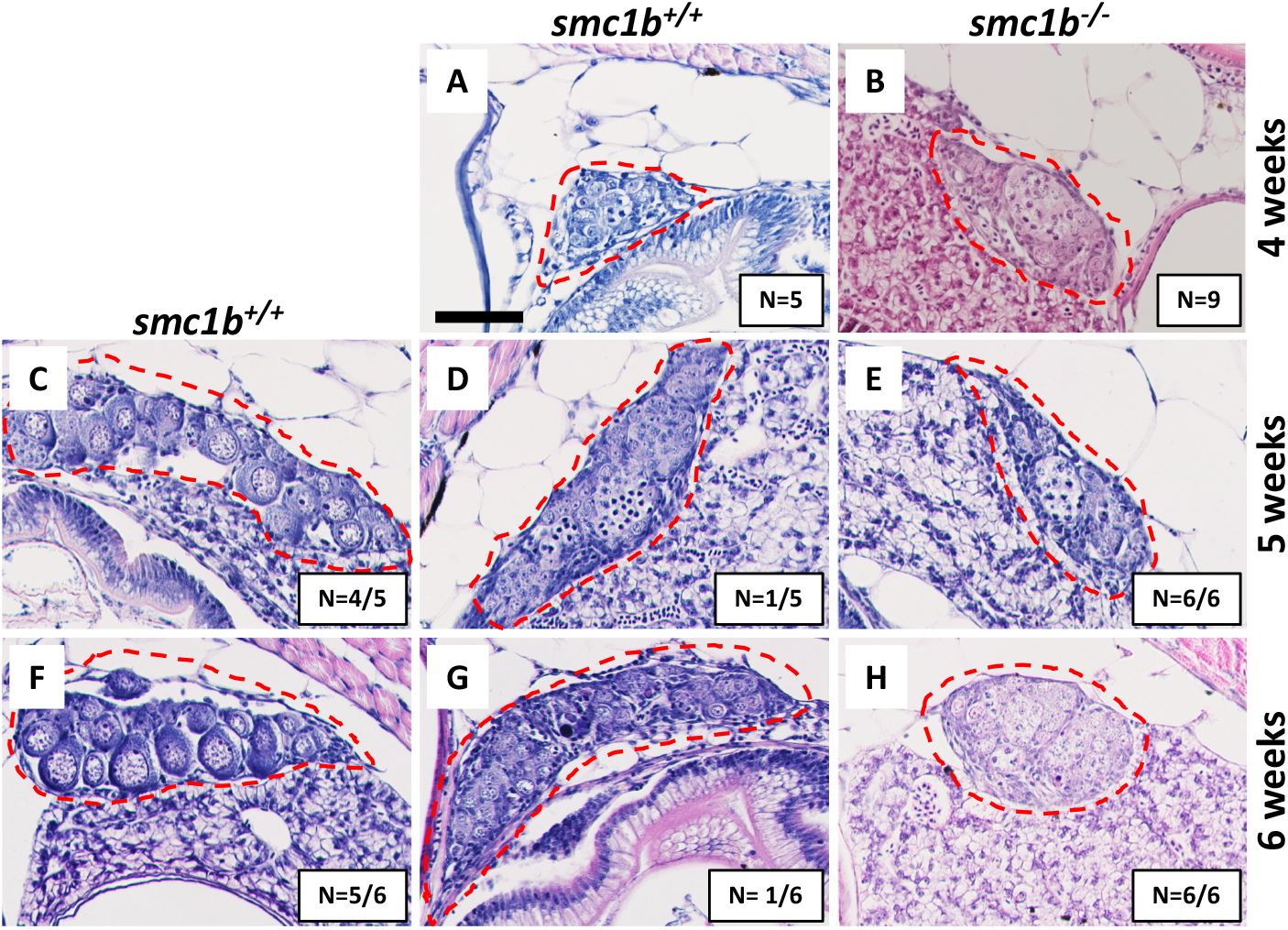
Absence of oogenesis in *smc1b^Q198X^* mutants. Hematoxylin-eosin stain of wild-type and *smc1b* mutant gonads during sex-differentiation. (A-B) Undifferentiated gonads in wild-type (A) and mutant (B) fish at 4 weeks old. (C-H) Although oogenesis (C, F) and spermatogenesis (D, G) took place in wild-type gonads at 5- and 6-weeks old, only spermatogenesis progressed in the mutants (E, H). Gonads are outlined by red dashed line, scale is 50 μm. N = total number of individuals with the gonad histology pictured per total number of fish analyzed for each age group.

### 3.3 *Smc1b* is necessary for meiotic bouquet formation and pairing and synapsis of homologous chromosomes

Since *smc1b* mutant testes had spermatocytes but no mature sperm (Fig 2C-E), we wanted to know when the spermatocytes arrested in meiosis. We stained mutant and wild-type testis sections with Sycp3 antibody and telomere fluorescent *in situ* hybridization (Telo-FISH) to determine the stage of meiosis in spermatocytes. In wild-type zebrafish, the telomeres attach to the nuclear envelope then cluster at one side of the nucleus to form the bouquet (Saito et al., 2014; Blokhina et al., 2019). Sycp3 begins to associate with the chromosomes in late leptotene to early zygotene stages, starting near the telomeres, then extends along the chromosomes as homologous chromosomes pair (Saito et al., 2014; Blokhina et al., 2018). We found that Sycp3-positive cells were present in the *smc1b* mutants, indicating that meiosis initiated in mutant testes (Fig 4E,G). However, in mutant spermatocytes, Sycp3 localization was comparable to wild type leptotene stage cells and spermatocytes resembling subsequent stages were not present (Fig 4A,C). Telomere dynamics were also affected in *smc1b* mutants. In wild type, telomeres begin clustering in late leptotene and form a tight cluster in early zygotene, forming the meiotic bouquet. We found some mutant spermatocytes with clustered telomeres, however these appeared less tightly clustered than wild-type bouquet-stage spermatocytes (Fig 4B,F). These data suggest that *smc1b* is necessary for progression from leptonema to zygonema.

**Figure 4:**
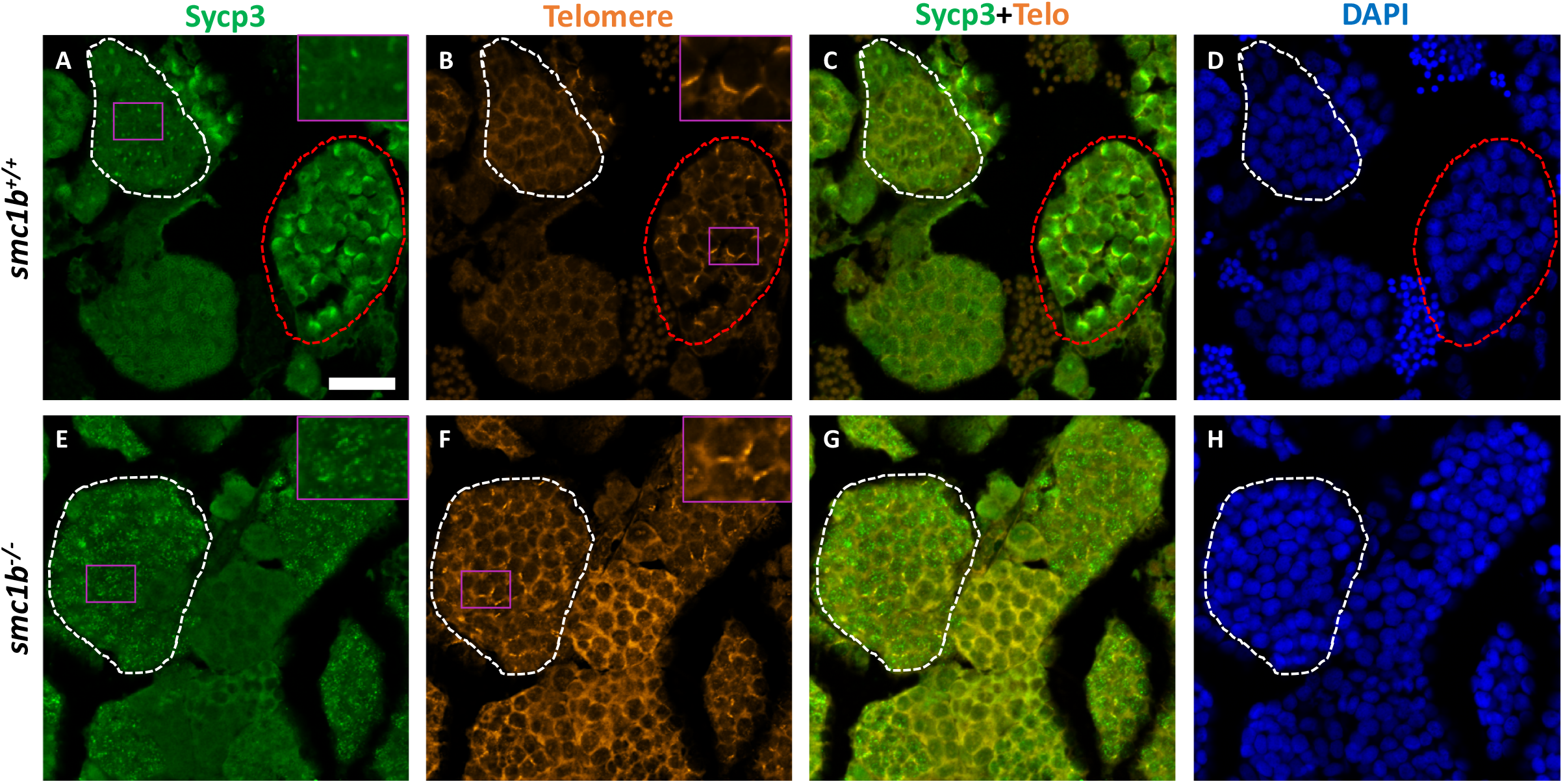
Spermatocytes in *smc1b^Q198X^* mutants failed to progress beyond leptotene stage. Labelling of Sycp3 and telomeres in adult wild-type and mutant testes sections: Sycp3 (green), telomere *in situ* hybridization (orange), DAPI nuclear stain (blue). (A-H) Leptotene stage spermatocytes are present in both wild-type and mutant spermatocytes, examples circled in white. Bouquet stage cells (circled red) are present in wild-type but not in the mutant. Insets are zoomed images of the areas boxed in purple. Scale bar is 20 μm.

To assess the stage at which *smc1b* mutant spermatocytes arrest in more depth, we performed Telo-FISH combined with IF for Sycp1 and Sycp3 antibodies on chromosome spreads (Fig 5). In wild type, Sycp1 protein associated with the synaptonemal complex beginning in early zygotene stage, as previously reported (Fig 5 A-D) (Saito et al., 2014; Blokhina et al., 2019). Similar to what we observed on sectioned testes, in *smc1b* mutant spermatocytes, Sycp3 only associated at the ends of chromosomes, resembling the leptotene stage in wild types (Fig 5E-F). Mutant spermatocytes had no Sycp1 protein associated with chromosomes, indicating that chromosomes failed to synapse in *smc1b* mutants (Fig 5E-F). Interestingly, we found about 5% of mutant spermatocytes exhibited some telomere clustering, which we refer to as “bouquet like” (Fig 5F). However, no mutant spermatocytes had tightly clustered telomeres characteristic of a fully formed bouquet, and Sycp1 labeling of the synaptonemal complex was absent. We therefore classified these “bouquet-like” spermatocytes as leptotene stage cells (Fig 5F-G). Overall, *smc1b* mutant spermatocytes failed to initiate synapsis between homologous chromosomes and progress to the zygotene stage, whereas 41% of wild-type primary spermatocytes were in the zygotene stage (Fig 5G). These results indicate that Smc1b is required for bouquet formation as well as pairing and synapsis of homologous chromosomes in zebrafish spermatocytes.

**Figure 5:**
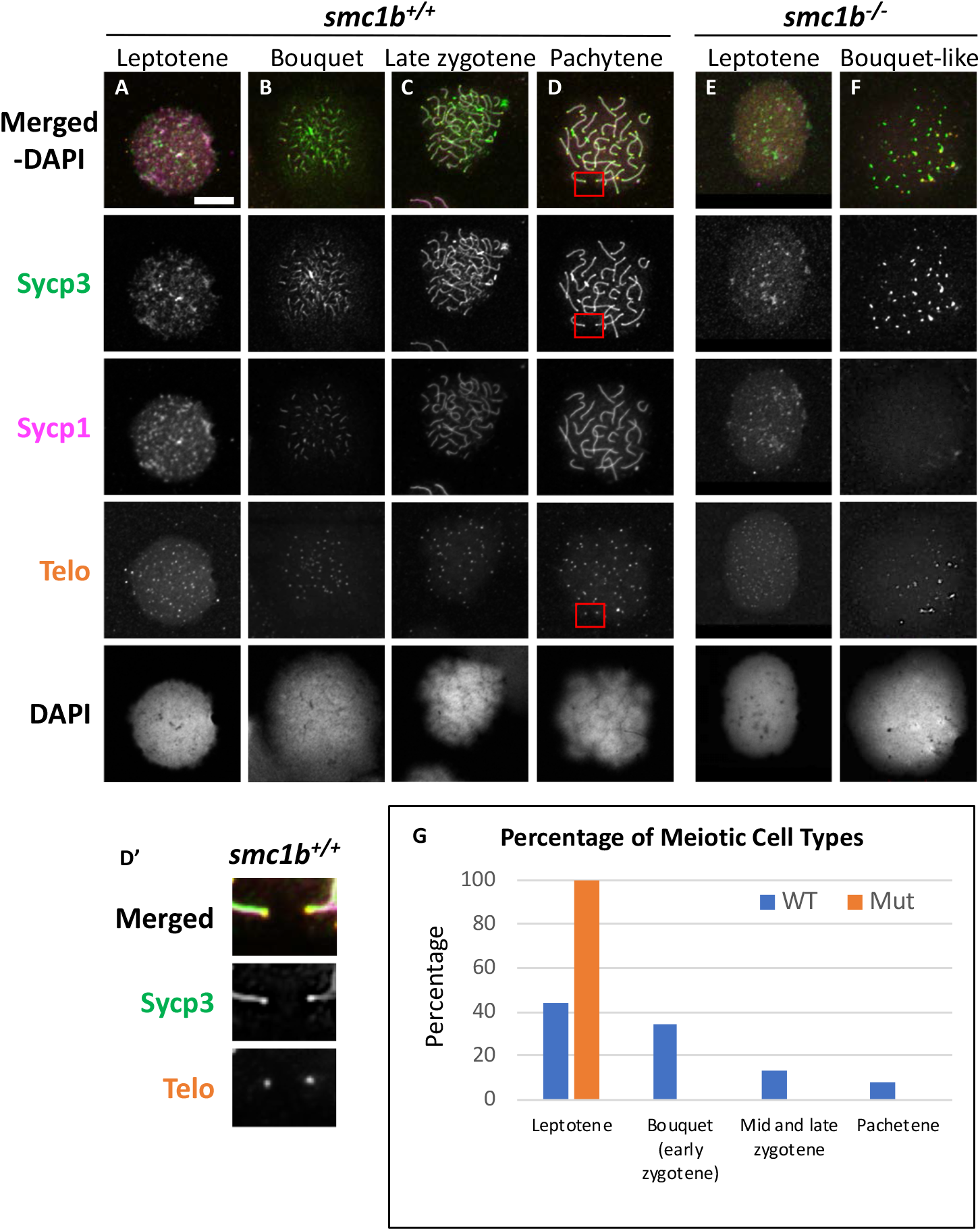
Failed pairing and synapsis in *smc1b^Q198X^* mutant spermatocytes. (A-F) Labelling of Sycp3 and Sycp1 proteins and telomeres (telo) on meiotic chromosome spreads from adult testes. (A-D) In wild-type spermatocytes, leptotene stage cells began expressing Sycp1 and Sycp3 protein. The synaptonemal complex proteins associated with chromosomes and assembled between homologous pairs in zygotene through pachytene stages as homologue pairing and synapsis occurred. (D’) Sycp3 exhibited enrichment at or near telomeres. Images show enlarged views of the red boxed areas in D. (E-F) *smc1b* mutant spermatocytes expressed Sycp1 and Sycp3 in leptotene stage, similar to wild type. However, Sycp3 only labeled chromosomes near the ends and Sycp1 did not associate with chromosomes. A small percentage of mutant cells exhibited some telomere clustering one side of the nucleus (F), which we called bouquet-like and classified as leptotene stage. (G) Quantification of spermatocytes showed that all mutant cells are in leptotene stage compared to about 40% in wild type. Later stages were not observed in the mutants (N = 52 wild-type and 130 mutant cells). Scale bar is 10 μm.

To ask if loss of Smc1b affects association of the cohesion complex with meiotic chromosomes, we assayed localization of the cohesion complex protein Smc3 (Fig 6). In wild type spermatocytes, Smc3 localization resembled to that of Sycp3 (Figs 5A-D, 6A-D). Smc3 associated with the synaptonemal complex in the bouquet stage starting near the chromosome ends (Fig 6B). By pachytene stage, Smc3 was localized along the entire synaptonemal complex (Fig 6D). Unlike what has been reported in mouse spermatocytes, Smc3 did not appear to be enriched near the telomeres in zebrafish, whereas Sycp3 was enriched (Fig 5D’, Fig 6D’) (Adelfalk et al., 2009). In the *smc1b* mutant leptotene stage cells, Smc3 protein was detected in nuclei but was not yet associated with chromosomes, similar to that of wild type (Fig 6A,E). However, while Smc3 associated with chromosomes at the bouquet stage of wild-type spermatocytes, Smc3 did not associate with chromosome axes in bouquet-like mutant spermatocytes (Fig 6F). These observations demonstrate that Smc1b is essential for cohesion complex formation during zebrafish meiosis.

**Figure 6:**
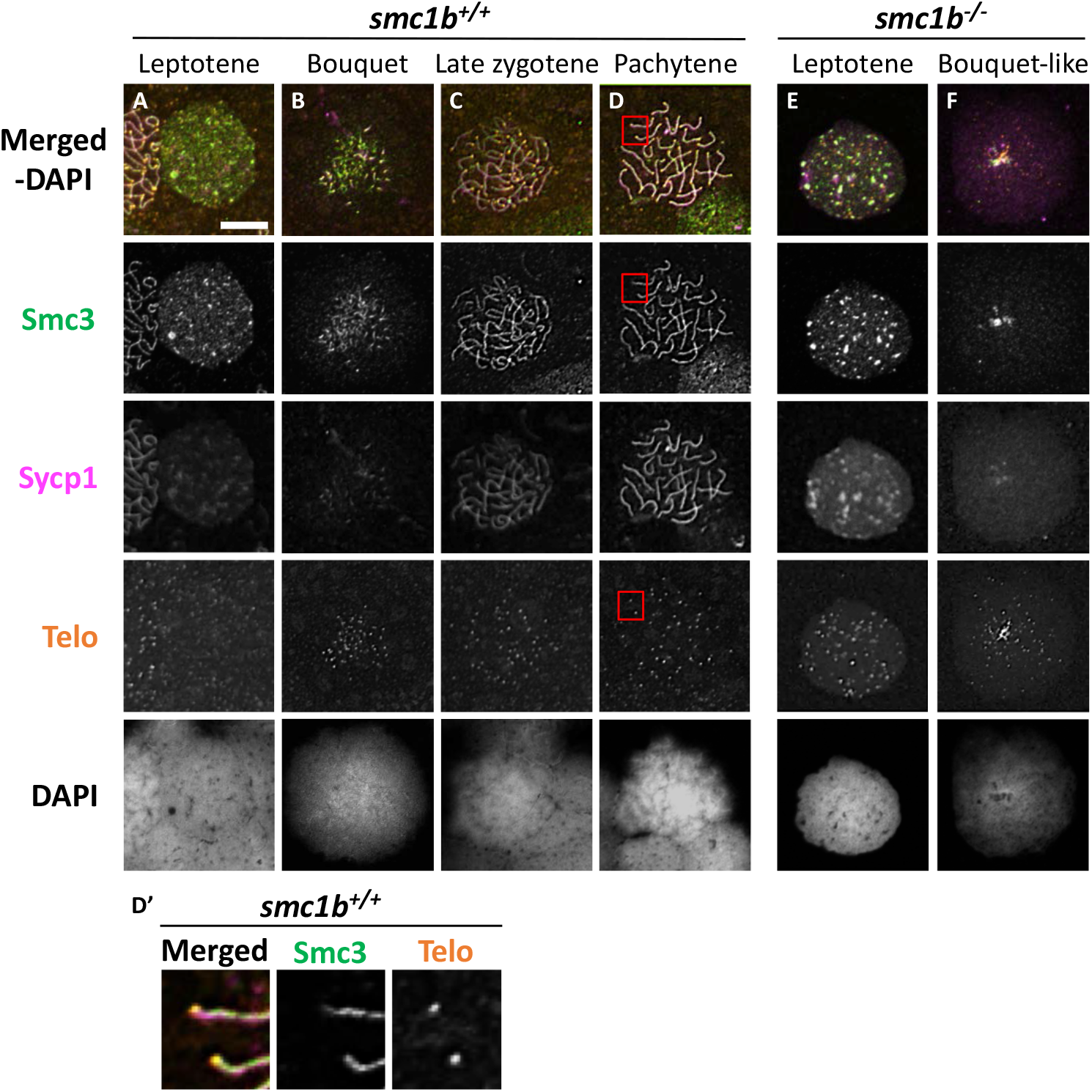
The cohesion complex does not associate with meiotic chromosomes in zebrafish spermatocytes. (A-E) Immunolabelling of Smc3, and Sycp1 in wild-type (A-D) and *smc1b* mutant (E-F) chromosome spreads. (A-D) In wild-type cells, Smc3 showed a localization pattern similar to Sycp1. (D’) Smc3 was not enriched at telomeres. Images show enlarged views of the red-boxed areas in D. (E-F) In mutant cells, Smc3 was expressed in leptotene stage but failed to load onto chromosomes. Scale is 10 μm.

### 3.4 Meiotic double strand breaks form in the absence of Smc1b

Formation of double strand breaks (DSB) is a pre-requisite of DNA recombination during meiosis. In meiotic cells, several proteins including histone variant γ-H2AX and Rad51 bind to DSBs and mediate exchange of DNA between homologous chromosomes (Hamer et al., 2003; Howard-Till et al., 2011). Therefore, expression and localization of these proteins are common markers of DSBs. To test whether zebrafish *smc1b* is required for meiotic DSB formation, we first performed IF on *smc1b* mutant testis sections with γ-H2AX antibody. In wild-type spermatocytes, γ-H2AX protein was initially localized near telomeres, as has been previously reported, which is most apparent in bouquet-stage cells in sectioned tissue (Fig 7A-D) (Saito et al., 2011; Blokhina et al., 2019). In tissue sections, early bouquet-stage cells, which are in late leptonema, appeared to have γ-H2AX localized more broadly within nuclei (Fig 7 A,D). In early zygotene stage, the telomeres became more concentrated on one side of the cell to form the late-bouquet. In these late-bouquet stage spermatocytes, γ-H2AX localization was also condensed on one side of the nucleus (Fig 7A,D). In *smc1b* mutant spermatocytes, γ-H2AX was detected but was not visibly concentrated to one side of the nuclei of spermatocytes (Fig 7E,H). Co-labeling with Telo-FISH, Sycp1, and γ-H2AX on nuclear spreads clearly demonstrated that γ-H2AX localization initiated near chromosome ends in late-leptotene and clustered near telomeres in the zygotene bouquet stage in wild-type spermatocytes (Supplementary Fig 2A-B). Similar to our observations of testes sections, in *smc1b* mutant spermatocytes, γ-H2AX was present but generally not concentrated to one side of the nucleus, resembling leptotene stage cells (Supplementary Fig 2E-F). These data indicate that DSBs could form in *smc1b* mutants despite lack of cohesion and synaptonemal complexes.

**Figure 7:**
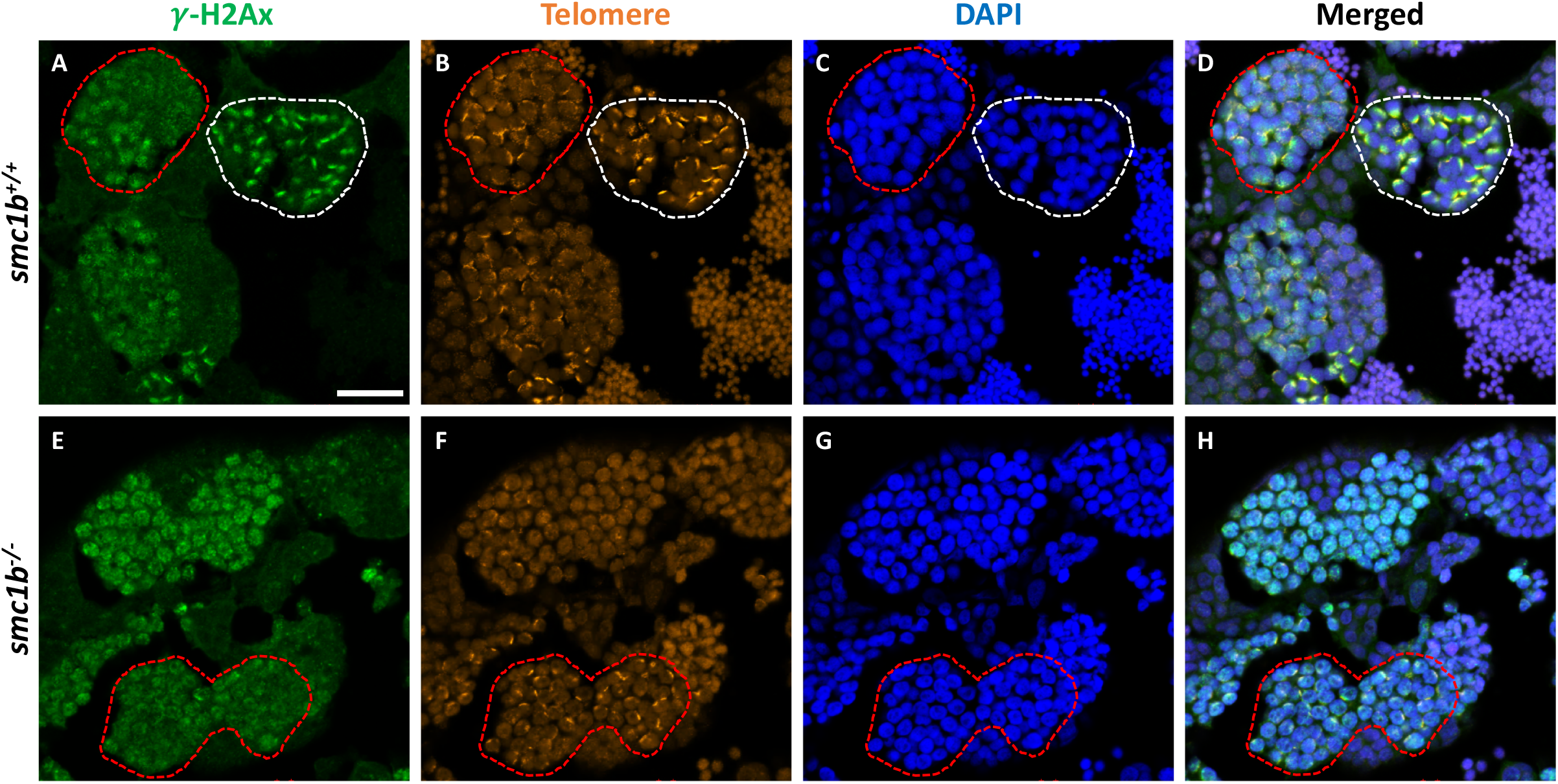
DNA double strand break marker γ-H2Ax was expressed in *smc1b^Q198X^* mutant testes. Immunolabeling of γ-H2Ax (green) and Telo-FISH (orange) on wild-type and mutant testis sections. (A-D) In wild-type early bouquet stage spermatocytes, γ-H2Ax appeared to localize more broadly in nuclei on testis sections (red dashed lines). Bouquet stage spermatocytes with tightly clustered telomeres (white dashed lines) exhibited γ-H2Ax localization near telomeres. (E-H) Early bouquet-stage spermatocytes in *smc1b* mutants display broad γ-H2Ax expression similar to wild type, however tight clustering of telomeres and γ-H2Ax to one side of the nucleus was not observed. Scale bar is 20 μm.

Finally, we stained chromosome spreads with Rad51 antibody to further assay the initiation of DSB repair of meiotic chromosomes in *smc1b* mutant spermatocytes. Following DSB formation, 5’-3’ end resection results in formation of single-stranded DNA that is bound by Dmc1 and Rad51. These proteins function to initiate DSB repair and strand invasion during meiotic recombination (Crickard and Greene, 2018). In wild-type spermatocytes, Rad51 protein was first localized near the chromosome ends in the bouquet-stage and then resolved into one or two spots per chromosome as early recombination nodules formed, as previously reported (Fig 8A-D) (Blokhina et al., 2019). In *smc1b* mutant spermatocytes, Rad51 was present in nuclei indicating that early DSB repair machinery necessary for meiotic recombination was active. Therefore, the meiotic cohesion complex was not required for initiation of meiotic DSBs in zebrafish spermatocytes.

**Figure 8:**
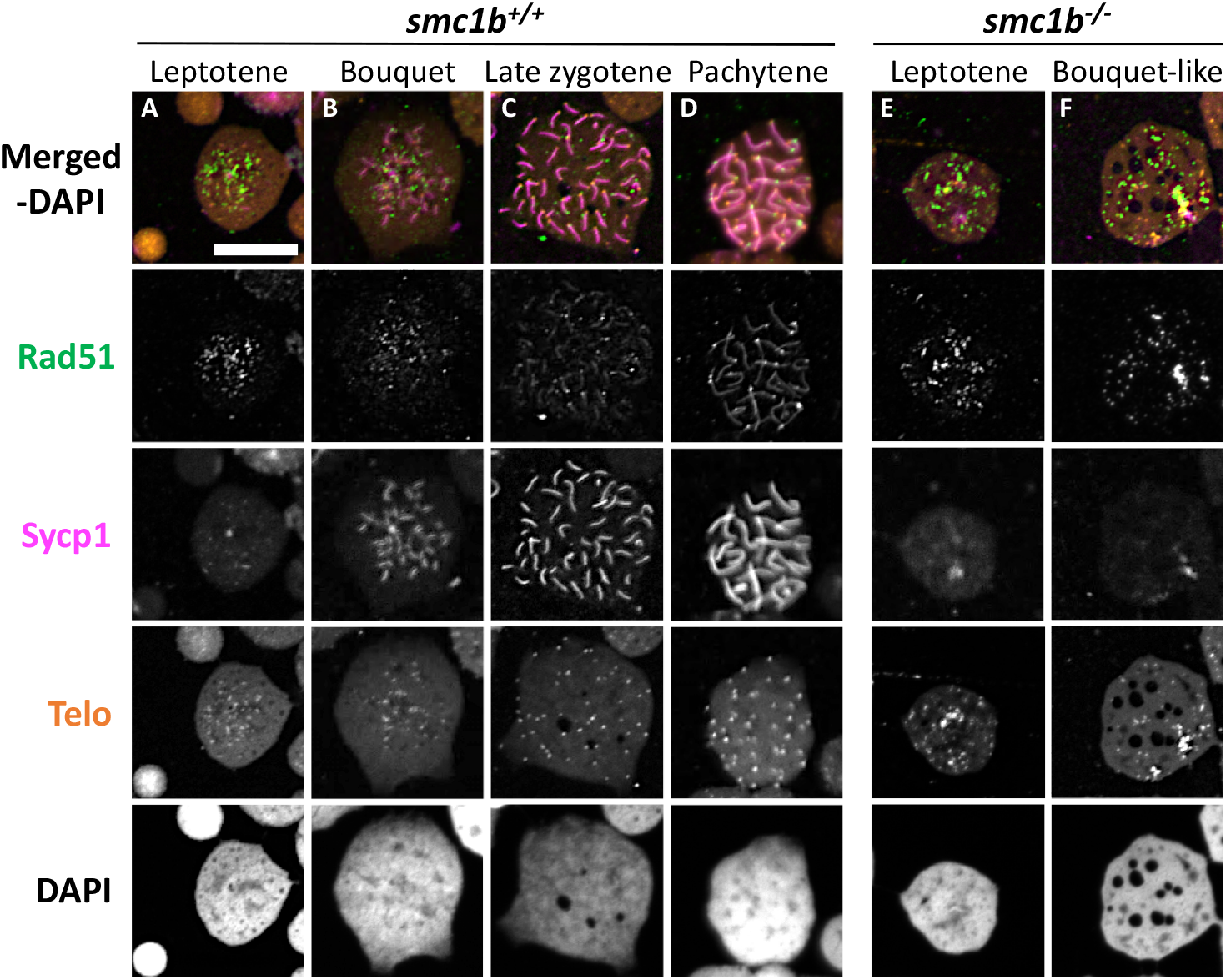
DNA double strand break and recombination marker Rad51 was expressed in *smc1b^Q198X^* mutant spermatocytes. Immunolabelling of Rad51 (green) and Sycp1 (magenta) in wild-type and mutant spreads, combined with telomere (telo) *in situ* hybridization (orange). (A-B) Rad51 protein was detected in wild-type beginning in leptotene stage and was initially located near the chromosome ends in leptotene and bouquet stages. (C,D) In more advanced wild-type spermatocytes, Rad51 localization become more distinct and resolved to one or two spots per chromosome. (E-F) Mutant spermatocytes had detectable levels of Rad51. Scale bar is 10 μm.

## 4 DISCUSSION

Vertebrate animals harbor two *smc1* genes, *smc1a* and *smc1β* (named *smc1a* and *smc1b* in zebrafish) In mammals, SMC1α functions primarily in mitotic divisions, whereas SMC1β predominantly functions in meiosis, although SMC1α has partially overlapping roles with SMC1β in mammalian meiosis (Biswas et al., 2018). The subfunctionalization of these two proteins as mitotic versus meiotic regulators has not been investigated outside of mammals. In this study, we demonstrate that the meiotic cohesion complex protein Smc1b is necessary for meiosis and fertility in zebrafish. *Smc1b* mutant zebrafish developed as sterile males without any other apparent defects. Histology showed that *smc1b* mutant spermatocytes entered meiosis but failed to complete the process as there was no sperm. We found that *smc1b* mutant spermatocytes arrested at the leptotene stage with a complete failure of homolog pairing and synapsis in the mutant cells. Nonetheless, meiotic DSBs still happened in *smc1b* mutant spermatocytes. Overall, our results indicate that Smc1b is essential for meiotic homologous chromosome pairing and synapsis in zebrafish, demonstrating that Smc1b functions as a meiotic cohesion protein in non-mammalian vertebrates, as has been found in mammals.

### 4.1 Differences in Smc1b requirement among vertebrates

We have shown that *smc1b* mutant zebrafish are infertile because of a failure to complete meiosis and an inability to produce mature sperm. Our finding is in agreement with previous studies in mammals, which showed that *smc1β* mutant male mice were sterile due to lack of spermatids and spermatozoa (Revenkova et al., 2004; Takabayashi et al., 2009). However, zebrafish *smc1b* mutants displayed more severe phenotypes compared to what has been reported in mouse mutants. For example, zebrafish *smc1b^-/-^* spermatocytes arrested at leptotene stage whereas mouse mutant spermatocytes were arrested in pachytene stage and oocytes in metaphase II stage. We found that zebrafish *smc1b^-/-^* spermatocytes failed to complete meiotic bouquet formation (early zygonema), even though some cells initiated telomere clustering. Furthermore, pairing and synapsis of homologous chromosomes did not occur in zebrafish *smc1b^-/-^* spermatocytes. By contrast, mice lacking SMC1β were able to form a synaptonemal complex (SC) between homologous chromosomes, although it was shorter than in the wild type and showed defects in late recombination markers but not early markers (Revenkova et al., 2004). Zebrafish *smc1b^-/-^* spermatocytes arrested in the leptotene stage, which is prior to when recombination takes place. However, we found that meiotic DSBs, which are a pre-requisite of recombination, formed in mutant spermatocytes and are therefore independent of *smc1b*.

One possible explanation for the discrepancies between the onset of the phenotype in zebrafish and mouse mutants is potential differences in Smc1a protein expression during meiosis. In mice, cohesion complex proteins, including SMC1α and SMC3, are localized to the homologous chromosome pairs in *Smc1β* mutant spermatocytes, albeit at reduced levels compared to wild type (Revenkova et al., 2004). Furthermore, expression of SMC1α in meiosis-I can partly substitute for SMC1β (Biswas et al., 2018). Zebrafish *smc1b* mutants may have earlier and more severe defects compared to mouse mutants due to a probable lack of Smc1a expression in spermatocytes. We attempted to label Smc1a protein in zebrafish to ask if Smc1a expression was absent in zebrafish spermatocytes, however the commercial antibodies we tested (abcam137707) did not work in zebrafish. Smc3 labeling demonstrated that zebrafish Smc3 did not localize to chromosomes in *smc1b^-/-^* spermatocytes, indicating that Smc1b is essential for formation of the meiotic cohesion complex in zebrafish. Interestingly, mutations that disrupt the mouse meiotic cohesion complex have similar defects to the zebrafish *Smc1b* mutants. For example, mice that were double mutant for *Rec8* and *Rad21L* or single mutant for *Stag3* failed to form axial elements or SCs and arrested in leptonema (Llano et al., 2012; Winters et al., 2014). These data suggest that, in contrast to mice, Smc1b is the only functional Smc1 cohesion subunit in meiotic germ cells in zebrafish.

We also found that there are some differences in *Smc1b* gene expression between zebrafish and mice in non-gonadal tissues. Previous studies reported that mouse SMC1β protein was expressed in the brain, heart, and spleen in addition to gonads (Mannini et al., 2015), but our result showed it primarily expressed in the gonads (Fig 1). In medaka, Smc1b protein was also detected specifically in the ovary and testis, however the only non-gonadal tissues assayed were the liver and OL32 cells (derived from fin tissue) (Iwai et al., 2004). Further exploration of differences in Smc1b expression across different vertebrate species would be informative in understanding the subfunctionalization of Smc1a and Smc1b between mitotically and meiotically dividing cell types in vertebrates.

We found that Smc3 colocalized with the synaptonemal complex in zebrafish, showing a similar localization pattern to that of mouse spermatocytes (Revenkova et al., 2001). However, in contrast to mice, we did not see Smc3 enrichment near telomeres in wild-type zebrafish spermatocytes (Fig 6). In mice, SMC1β, SMC3, STAG3, and SYCP2 have all been reported to be enriched near the telomeres in meiotic prophase-I (Liebe et al., 2004; Adelfalk et al., 2009). Interestingly, we detected enrichment of Sycp3 near the telomeres, but did not detect enrichment of Smc3 in zebrafish (Figs 5,6). Cohesion complex enrichment at or near telomeres is thought to maintain telomere integrity and telomere attachment to the nuclear envelope during meiosis – *Smc1β* mutant spermatocytes exhibited loss of SMC3 telomere enrichment and had defects in telomere attachment and integrity in mice (Adelfalk et al., 2009). The presence of enriched Sycp3 instead of Smc3 in zebrafish indicates that there could be different mechanisms of telomere attachment and maintenance during meiosis in these two species.

### 4.2 Crosstalk between cohesion, the synaptonemal complex, and DSBs during zebrafish meiosis

The reason for failed homolog pairing and synapsis in zebrafish Smc1b mutants could be impaired interactions between cohesion complex proteins and SC proteins. The vertebrate SC consists of axial element proteins Sycp2 and Sycp3, transverse filament protein Sycp1, and multiple central element proteins (Gao and Colaiácovo, 2018). In mice, cohesion subunits SMC1 and SMC3 physically interact with SC axial element proteins SYCP2 and SYCP3 (Yang and Wang, 2009). In zebrafish spermatocytes, Smc1b is required for Sycp3 and Sycp1 to associate along chromosomes suggesting that interactions between these proteins is necessary for SC formation. This notion is further supported by recent analysis of zebrafish SC mutants, which also show defects in homolog pairing and synapsis (Takemoto et al., 2020; Imai et al., 2021). For instance, spermatocytes mutant for *sycp2,* encoding an SC axial element protein, failed to form an SC, which disrupted homologue pairing (Takemoto et al., 2020). Interestingly, these mutants did not have the axial element protein Sycp3 associated with chromosomes whereas the transverse filament protein Sycp1 did associate with chromosomes. In the *sycp2* mutants, the meiotic cohesion complex protein Rad21l1 colocalized with Sycp1 on chromosomes (Takemoto et al., 2020). We found that Smc1b was necessary for the cohesion complex (Smc3), Sycp3, and Sycp1 to associate with chromosomes in meiosis. Together, these data suggest that the cohesion complex may be involved in recruiting Sycp1 to chromosomes, even in the absence of axial element proteins. Mutations disrupting the SC transverse element protein Sycp1 displayed somewhat less severe defects than those observed for *sycp2* and *smc1b* mutants. Homologous chromosomes initially paired in Sycp1 mutant spermatocytes, however synapsis did not occur which caused loss of pairing and germ cell arrest at late zygotene/early pachytene stage (Imai et al., 2021). Analysis of these mutants demonstrates that the zebrafish cohesion complex is intimately associated with SC formation and function during meiosis, as has been observed in other organisms.

Smc1b was not required for DSBs suggesting that a functional meiotic cohesion complex is not a prerequisite for meiotic DSBs in zebrafish spermatocytes. DSB formation during meiosis is induced by topoisomerase-type enzyme Spo11 (Keeney et al., 1997). Zebrafish lacking Spo11 formed the meiotic bouquet despite failed DSB formation, however homolog pairing and synapsis were disrupted in the mutant spermatocytes (Blokhina et al., 2019). Formation of meiotic DSBs was also disrupted in *sycp2* mutants. These mutants also displayed failure of the axial element protein, Sycp3, to associate with chromosomes (Takemoto et al., 2020). By contrast, we found that *smc1b* mutant spermatocytes lacked formation of the chromosome axes but could still form DSBs. Based on these observations, we propose that failure to form chromosome axes does not preclude meiotic DSBs, however axial element proteins may facilitate Spo11-mediated DSB formation. Despite DSB formation, *smc1b* mutants spermatocytes arrested in late-leptotene stage with failed pairing and synapsis. Therefore, it is unlikely that DSBs that formed in zebrafish *smc1b^-/-^* spermatocytes would resolve into recombination events. These findings are consistent with mouse mutants lacking a meiotic cohesion complex (e.g. *Stag3^-/-^* or *Rad21/^-/-^ Rec8^-/-^*), where DSBs occurred despite a failure to form chromosome axial elements and a synaptonemal complex (Llano et al., 2012; Winters et al., 2014).

### 4.3 Failure of female development in zebrafish meiotic mutants

We found that all *smc1b* mutant zebrafish developed as male, demonstrating that *smc1b* is required for female development. In zebrafish, formation of ovarian follicles is a prerequisite for establishment and maintenance of the ovary (Kossack and Draper, 2019). Folliculogenesis involves the separation of oocytes, which are initially clustered together within nests, into individual oocytes each of which is surrounded by somatic cells. Generally, oocytes initiate meiosis while in the nests and arrest in meiosis I after follicles form (Elkouby and Mullins, 2017). In zebrafish, folliculogenesis begins when oocytes are in the pachytene stage and oocytes arrest in the diplotene stage as oocytes continue to grow and follicles develop (Elkouby et al., 2016). Meiosis-I resumes at the maturation stage, which occurs just prior to ovulation in zebrafish (stage IV), then arrests again in meiosis-II until fertilization takes place (Selman et al., 1993). Multiple meiotic mutants have all male phenotypes in zebrafish, including mutants disrupting *sypc1* and *sycp2* genes (Saito et al., 2011; Takemoto et al., 2020; Imai et al., 2021). In mutants affecting synaptonemal complex components, oogenesis was initiated, as juvenile fish had ovarian follicles at 40 dpf *(sycp1^-/-^*) and 28 dpf *(sycp2^-/-^*) (Takemoto et al., 2020; Imai et al., 2021). Therefore, in the *sycp1* and *sycp2* mutants, oogenesis did not proceed to stages that could support ovary development into adulthood. However, in both *spo11* and *mlh1* mutant zebrafish, fish developed into adult males and females (Feitsma et al., 2007; Blokhina et al., 2018). The *mlh1* mutant zebrafish could complete meiosis but showed recombination defects and produced aneuploid progeny (Feitsma et al., 2007; Leal et al., 2008). Thus, sex reversal is not universal in zebrafish meiotic mutants, but it is a common phenotype in mutants that fail to complete meiosis. Previous studies found that *smc1b* was expressed in bipotential goands of juvenile zebrafish, therefore it is likely to play an important role in meiosis in the juvenile ovary (Tzung et al., 2015). We found that ovarian follicles failed to form in *smc1b^-/-^* gonads. Because *smc1b* mutants did not form ovarian follicles in juvenile fish, it is likely that oocytes arrested prior to the pachytene stage.

---

**Supplementary Figure 1:** Conservation of key motifs in vertebrate Smc1b proteins. (A) Graphical representation of zebrafish Smc1b protein. Numbers inside boxes represent amino acid position in the corresponding domain. (B) Percentage of identical amino acid residues between human (h) and zebrafish (z), and mouse (m) and zebrafish Smc1b proteins. (C-E) Protein sequence alignments of the N-terminal ATP binding domain (C), flexible hinge domain (D), and C-terminal ATP binding domain (E). The conserved motifs are highlighted in green. The consensus motif sequences are Walker A: GxxGxGK(S/T), ABC transporter signature motif: LSGG(E/Q) (K/R), Walker B: hhhhDE, where x is any of 20 amino acid and h is any hydrophobic residue.

**Supplementary Figure 2:** DNA double strand break marker γ-H2Ax was expressed in *smc1b^Q198X^* mutant testes. (A-F) Labeling of γ-H2Ax (green), Sycp1 (magenta), and Telo-FISH (orange) on wildtype (A-D) and mutant (E-F) chromosome spreads from testes. γ-H2Ax protein is expressed in both wild-type and mutant nuclei. Scale bar is 10 μm.

## AUTHOR CONTRIBUTIONS

KNI contributed to data shown in all figures. MMM contributed to data shown in Figures 2 and 3. KNI and KRS designed the experiments, analysed data, and wrote the manuscript.

## ACKNOWLEDGMENTS

We are grateful to Sean Burgess and James Amatruda for providing Sycp1 and γ-H2Ax antibodies, respectively. We are also thankful to The Zebrafish Mutation Project for generating the *smc1b* mutants and to the Zebrafish International Resource Center (ZIRC) for supplying *smc1b* mutant zebrafish. Special thanks to Jessica MacNeil and Kavita Venkataramani for helpful discussions about the manuscript.

