## Supplemental Figure 1 for "The zebrafish meiotic cohesion complex protein Smc1b is required for key events in meiotic prophase I"

A

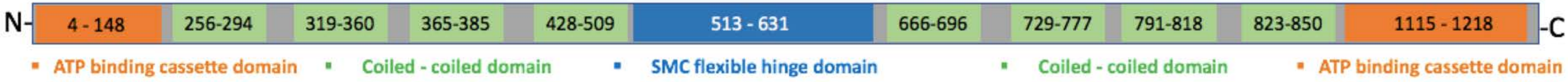


B

| Smc1b protein Domains | % identical sequence between zebrafish and: | |
|---|---|---|
| | Human | Mouse |
| Whole protein | 53.04 | 52.11 |
| N-terminal ATP binding domain | 62.75 | 64.13 |
| Coiled-coiled domain 1 | 56.41 | 48.72 |
| Coiled-coiled domain 2 | 28.57 | 35.71 |
| Coiled-coiled domain 3 | 57.14 | 57.14 |
| Coiled-coiled domain 4 | 52.44 | 50.00 |
| Flexible hinge domain | 66.95 | 64.41 |
| Coiled-coiled domain 5 | 35.48 | 35.48 |
| Coiled-coiled domain 6 | 30.61 | 26.53 |
| Coiled-coiled domain 7 | 53.57 | 53.57 |
| Coiled-coiled domain 8 | 32.14 | 35.71 |
| C-terminal ATP binding domain | 75.96 | 75.96 |

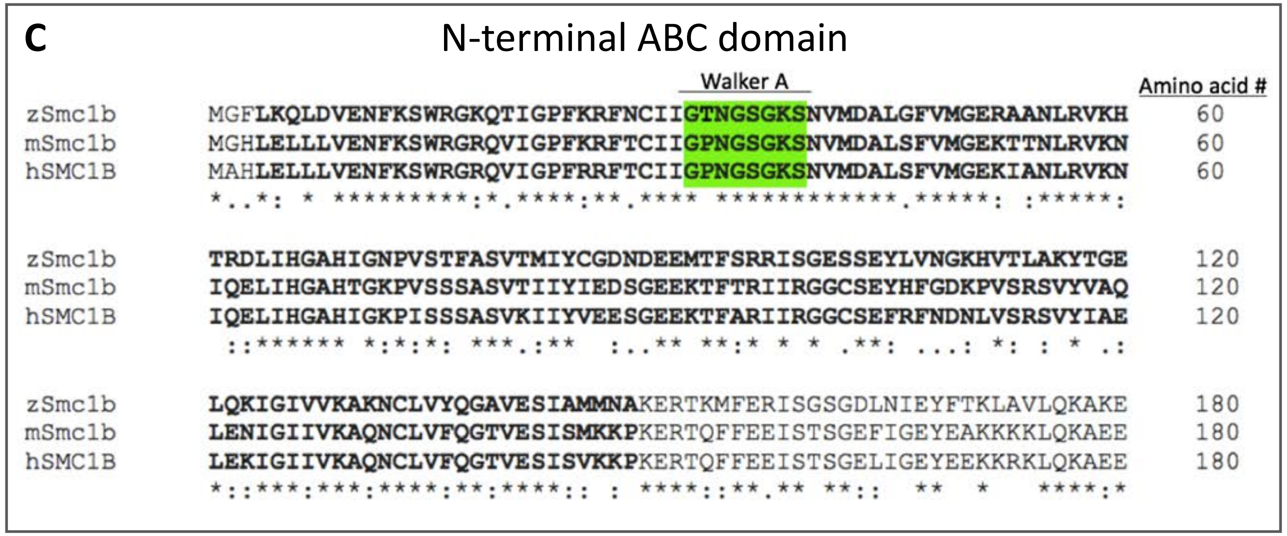


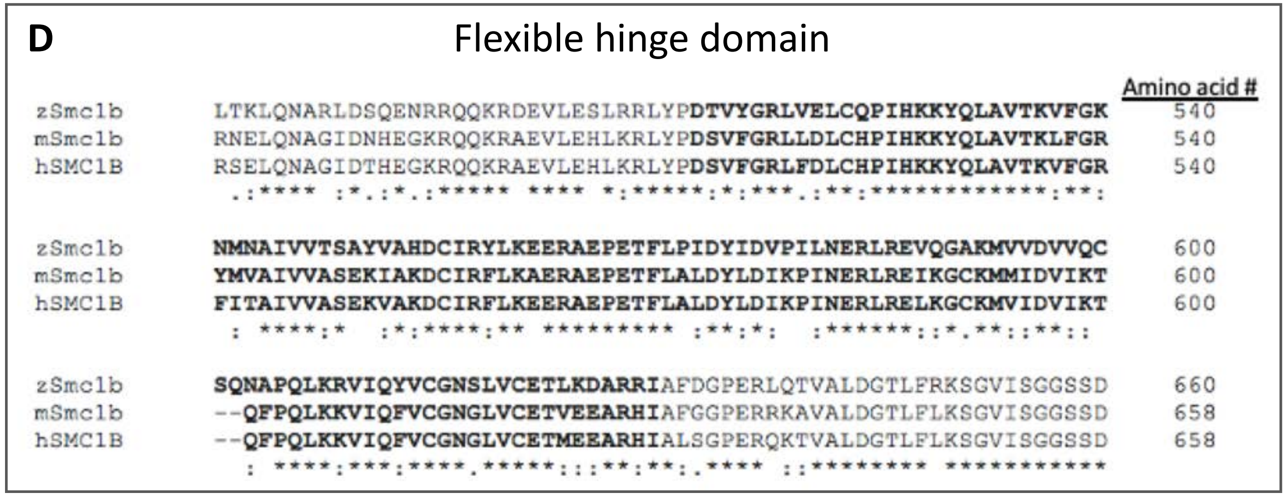


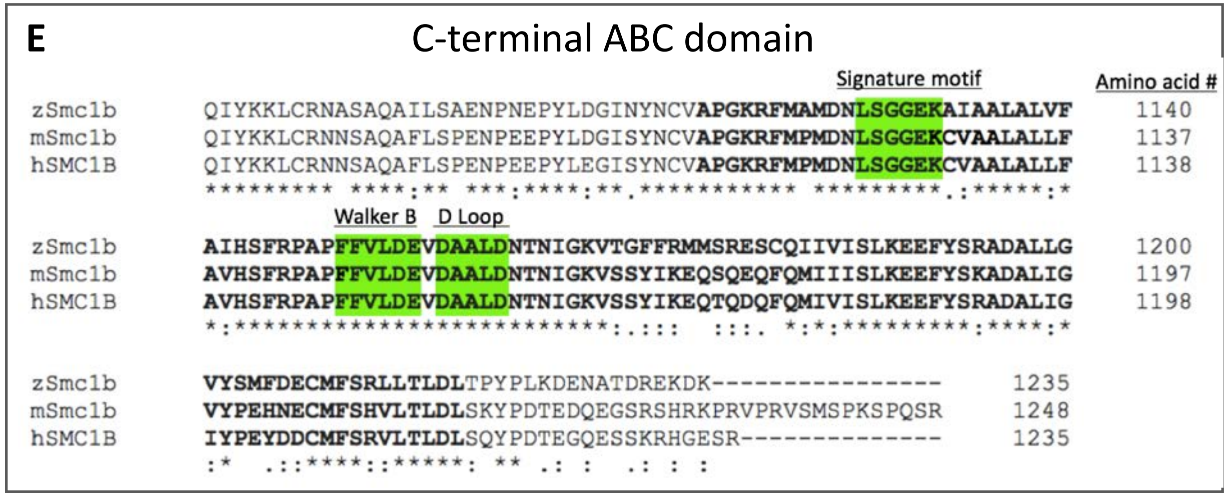
