## Supplementary figures and images for "The zebrafish meiotic cohesion complex protein Smc1b is required for key events in meiotic prophase I"

### Supplemental Figure 2

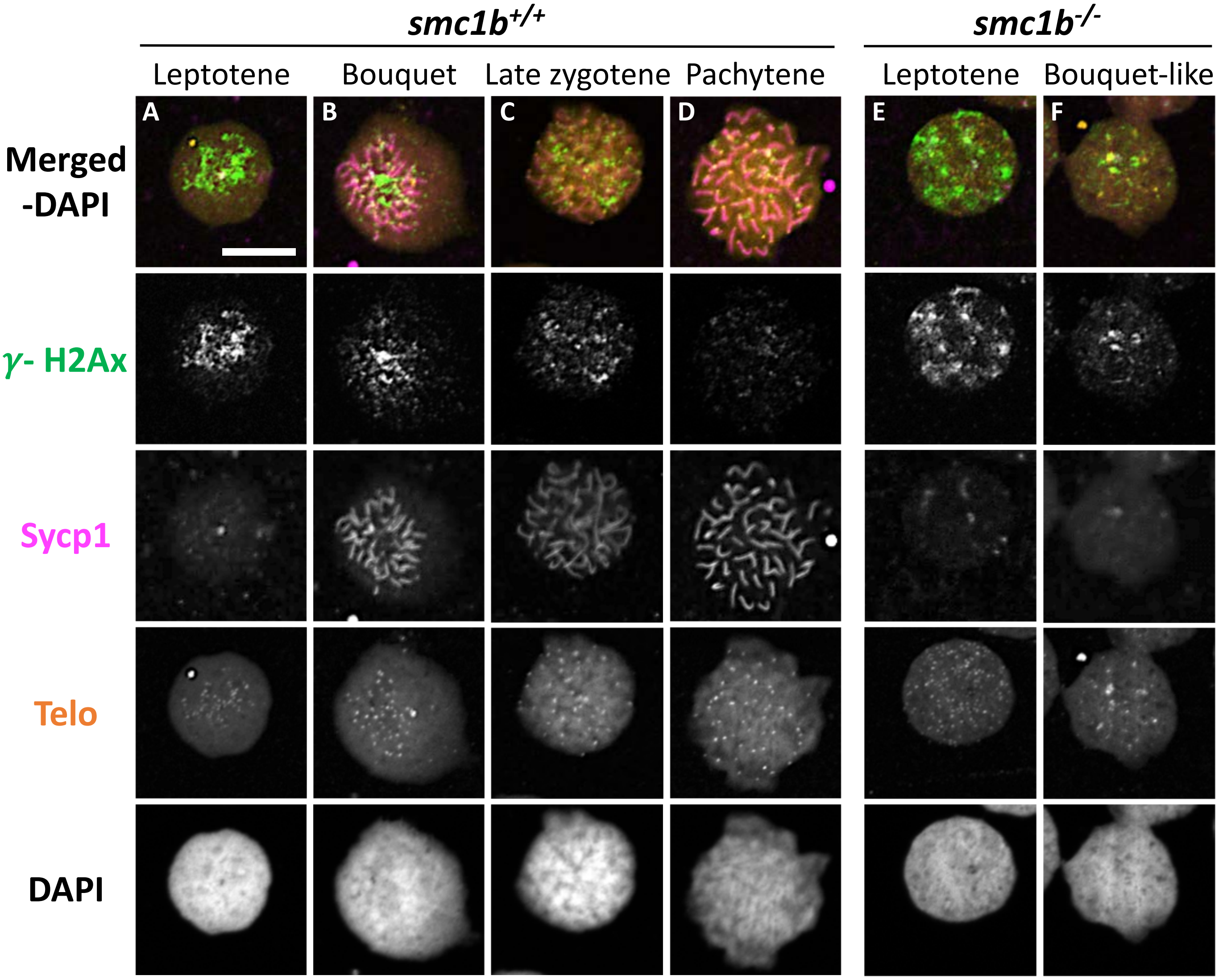
